# Motor outcomes congruent with intentions may sharpen metacognitive representations

**DOI:** 10.1101/2022.04.19.488801

**Authors:** Angeliki Charalampaki, Caroline Peters, Heiko Maurer, Lisa K. Maurer, Hermann Müller, Julius Verrel, Elisa Filevich

**Author notes:** **Corresponding Author:** Angeliki Charalampaki.

## Abstract

We can monitor our intentional movements, in order to describe how we move our bodies. But it is unclear which information this metacognitive monitoring relies on. For example, when throwing a ball to hit a target, we might use the visual information about how the ball flew to metacognitively assess our performance. Alternatively, we might disregard the ball trajectory — given that it is not directly relevant to our goal — and metacognitively assess our performance based solely on whether we reached the goal of hitting the target. In two experiments we aimed to distinguish between these two alternatives and asked whether the distal outcome of a goal-directed action (hitting or missing a target) informs the metacognitive representations of our own movements. Participants performed a semi-virtual task where they moved their arm to throw a virtual ball at a target. After each throw, participants discriminated which of two ball trajectories displayed on the screen corresponded to the flight path of their throw and then rated their confidence in this decision. The task included two conditions that differed on whether the distal outcome of the two trajectories shown matched (*congruent*) or differed (*incongruent*). Participants were significantly more accurate in discriminating between the two trajectories, and responded faster in the *incongruent* condition and, accordingly, were significantly more confident on these trials. Crucially, we found significant differences in metacognitive performance (measured as meta-d’/d’) between the two conditions only on successful trials, where the virtual ball had hit the target. These results indicate that participants successfully incorporated information about the outcome of the movement into both their discrimination and confidence responses. However, information about the outcome selectively sharpened the precision of confidence ratings only when the outcome of their throw matched their intention. We argue that these findings underline the separation between the different levels of information that may contribute to body monitoring, and we provide evidence that intentions might play a central role in metacognitive motor representations.

**Highlights:**

- Participants threw a virtual ball to hit a target
- Following their throw participants selected between two plausible trajectories
- On half the trials, the two alternative trajectories differed in their distal outcome.
- Discrimination and confidence were higher in trials with different outcome.
- Metacognitive performance was best in hit trials when the alternative missed.

## 1. Introduction

Moving our body seems to happen with precision and effortlessly, while we attend to the world around us. According to motor control theories, the motor system issues the commands necessary to transition from the current to the intended body position to achieve a goal (Blakemore et al., 2002). An efference copy of these commands is used to evaluate the accuracy of the movement performed and apply any necessary corrections (Miall & Wolpert, 1996; Wolpert et al., 1995; Wolpert & Flanagan, 2001), which can happen in the absence of awareness (Bourdin et al., 2019; Fourneret & Jeannerod, 1998; Gaveau et al., 2014; Slachevsky et al., 2001)

A given motor goal is not enough to specify the necessary motor commands, because virtually any goal-directed movement can be achieved through a manifold combination of muscular activity, following the principle of motor abundance (Latash, 2000). That is, any movement needs only satisfy the constraints that ensure that the goal is reached, but can, and does, vary over repetitions (Latash, 2012). Despite this variability in the low-level details of each movement, previous evidence suggests that the brain can accurately predict the movement outcome, i.e., whether the motor goal was reached or not. In particular, two studies employed a semi-virtual ball throwing task in which participants used a manipulandum to grab and throw a virtual ball to hit a target (Joch et al., 2017; Maurer et al., 2015). Participants completed the same motor task in both studies, but received different visual information (the ball trajectory and whether they hit the target or not in one case; and no feedback about the ball trajectory, but delayed feedback about whether they hit the target, in the other case). Both studies revealed that an error-related negativity in the electroencephalography (EEG) signal occurred right after making an erroneous movement (i.e., that led to a target miss). This suggests that outcome predictions occur early on, and independently from explicit visual feedback. Importantly, however, these outcome predictions are not free from error. A separate study in which participants completed the same motor task and verbally reported their predicted outcome, showed that participants often made mistaken predictions (Maurer et al 2022). Further, participants have been shown to misrepresent the outcome of their own movements (overestimating their performance) when they themselves pressed a button to stop a moving ball at a cued location, but were accurate when estimating the outcome in a control visual movement replay condition (Wolpe et al., 2014). Hence, because any given motor goal can be achieved in an abundance of ways, and because small body adjustments can happen during motor execution without conscious control, it follows that the low-level details of motor control might escape conscious monitoring. Instead, motor monitoring might focus on performance, and rely on (often noisy) information about the outcome of the movements. To test this hypothesis, we adapted a semi-virtual motor task where participants made ecologically valid, goal-oriented movements and threw a virtual ball to hit a virtual target (Joch et al., 2017; Maurer et al., 2015). In our study, after each ball throw, participants discriminated between two plausible trajectories (one real and one alternative) to indicate which one they thought corresponded to the ball flight trajectory following their throw, and then rated their confidence in their own responses. In two conditions, participants completed two types of trials, where the outcome of the alternative trajectory was either congruent or incongruent with the real one. Specifically, we manipulated the alternative ball trajectory to control whether it led to a successful distal outcome (hitting the target) or not (missing the target). To determine whether higher-order representations rely primarily on monitoring the outcome of a movement, as opposed to the lower-level details, we estimated metacognitive efficiency for each experimental condition, which quantifies the relationship between confidence judgments and accuracy in the discrimination task (Fleming & Lau, 2014). If motor outcome is indeed what motor monitoring is based on, we hypothesized that metacognitive efficiency would be higher on trials where the distal outcome differed between the two trajectories that participants chose from, as in these trials the distal outcome of the movement would be informative for the discrimination decision.

## 2. Methods

### 2.1 Experiment 1

The experiment was pre-registered (https://osf.io/v635y/), and we adhered to the pre-registered plan unless stated otherwise.

#### 2.1.1 Participants

Forty-six healthy, right-handed participants (26.4 ± 4.5 years old, 31 female) took part in the study. Handedness data, collected post-hoc from 21 participants using the Edinburgh Handedness Inventory, confirmed that participants were right-handed, (mean score ± SD: 91 ± 11). Participants reported no neurological or psychiatric history and normal or corrected-to-normal vision. They received detailed instructions in English or German, signed written informed consent prior to starting the experiment, and received 8 €/hour as compensation for their time. The study was conducted according to the Declaration of Helsinki and was approved by the ethics committee of the Institute of Psychology of the Humboldt-Universität zu Berlin.

#### 2.1.2 Apparatus and Stimuli

The motor task consisted of a virtual version of the “Skittles” game (Müller & Sternad, 2004; Sternad et al., 2011) programmed using Matlab (R2016b, MathWorks, Natick, MA) and Psychtoolbox-3 (Brainard, 1997; Kleiner et al., 2007; Pelli, 1997). In the Skittles game, participants swing a ball, attached with a rope from the top of a pole, aiming to hit a target — the skittle — that stands behind the pole (Figure 1.A). In the virtual version of the game, participants sat approximately 60 cm away from an LCD monitor (2560 × 1440 pixels, 61 × 34.5 cm, refresh rate 60 Hz) and rested their right hand on a custom-made lever, which could rotate on a vertical axis under the participant’s elbow, allowing them to bend and straighten their elbow on the horizontal plane. To record the angle of the lever (i.e., of participants’ elbows), we used a goniometer (Novotechnik, Stuttgart, Germany, RFC4800 Model 600, 12 bit resolution, 0.1° precision) placed on the rotation axis of the lever. Additionally, the lever had a touch-switch at the distal end. Participants “grabbed” the virtual ball by placing their index finger on the tip of the lever, and released it by lifting their finger. We recorded data from the lever using a Labjack T7 data acquisition device (LabJack Corp., Lakewood, CO) with a sampling rate of 1 kHz.

**Figure 1.**
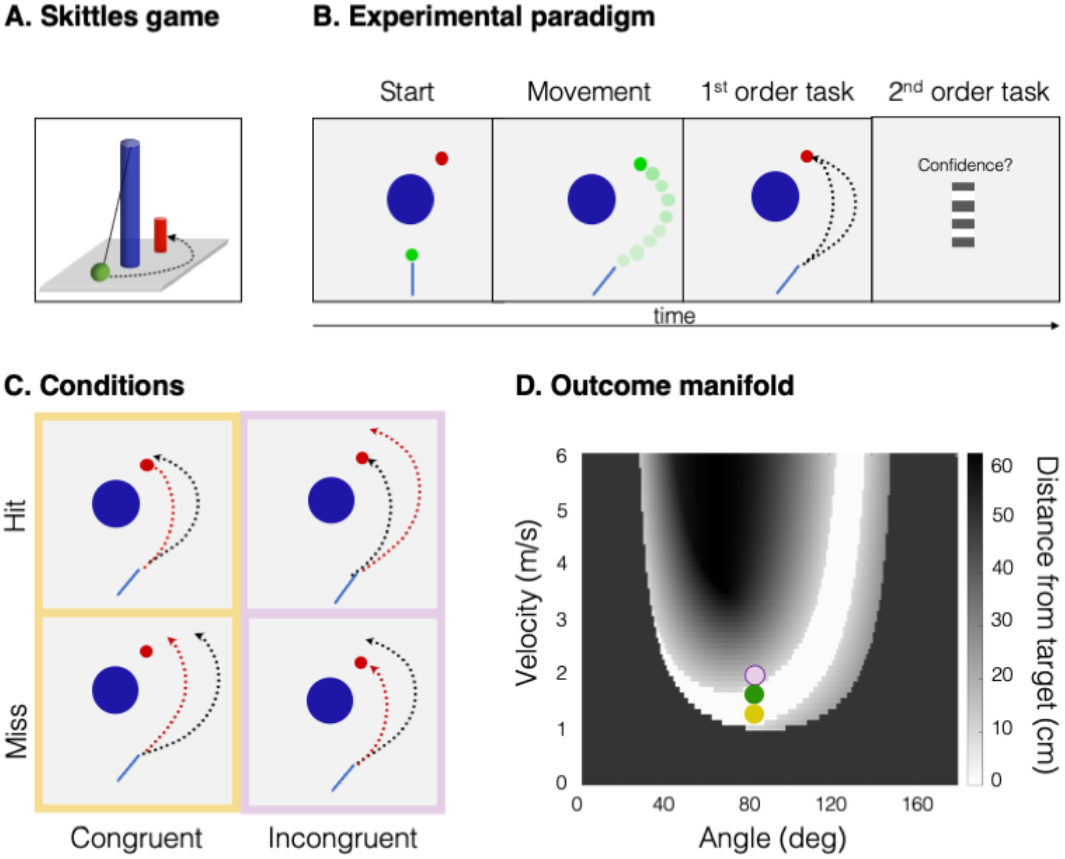
Experimental setup and paradigm. A. Sketch of the Skittles task in perspective showing the blue pole, green ball and red target. **B. Experimental paradigm**. On each trial participants threw a virtual ball in order to hit a target. After each throw, they discriminated which of the two displayed trajectories best corresponded to the movement they had just made. Finally, participants rated their confidence in the preceding discrimination decision. **C. Conditions**. The two conditions included in the experiment differed only in whether the alternative trajectory (red) had the same distal outcome as the actual trajectory or not (respectively, *congruent*/*incongruent* conditions, framed in yellow/pink). **D. Outcome manifold** for the Skittles task. The combination of ball release parameters (angle of release and velocity of the tip of the lever at the point of release) fully determines the trajectory of the ball, and therefore the minimum distance between ball and target). The regions shown in white correspond to combinations of release angles and velocities that result in hitting the target. The areas indicated with grayscale correspond to combinations that result in missing the target, while the black areas correspond to those that result in hitting the central pole. We illustrate with a green circle the real combination of angle and velocity of an example trial. By adding or subtracting the same value from the real velocity it is possible to draw an alternative trajectory that has incongruent (pink) or congruent (yellow) outcomes respectively.

#### 2.1.3 Procedure

The main experiment consisted of 480 trials, split into four blocks and took approximately 90 minutes. On the screen, participants saw a bird’s-eye view of the Skittles scene (Figure 1.B), which included the lever (represented as a bar that rotated around its end, along with the physical lever), the central pole (a central large blue circle), the target (red circle placed behind the central pole, to the right of the scene midline), and a ball, depicted in green. Participants started the trial by picking up the virtual ball: They placed their index finger on the sensor at the end of the lever and swung the virtual ball around the pole by extending their elbow and lifting their index finger to release the ball. We specified the Skittles model by setting the following constant values (for further details, see Müller & Sternad, 2004; Sternad et al., 2011): central pole radius = 0.25 m; central pole position (x,y) = (0 m,0 m); initial ball and target radius = 0.05 m (but see the section on Online staircases for details on how this changed according to participants’ behavior); target position (x,y) = (0.4 m, 0.5 m); massless rope constant k = 1 N/m. In this deterministic task, the ball flight trajectory is defined by two parameters only, namely the velocity of the tip of the lever and the angle of the lever at the point of release (Müller & Sternad, 2004; Sternad et al., 2011). The ball was then shown flying around the pole, returning to the vertical midline where the axis of the lever was shown. The target disappeared from the scene at the time of ball release, so participants did not receive any explicit information about whether they had hit the target.

After each ball throw, participants saw a static Skittles scene including the lever, pole, target and two lines representing sections of two plausible trajectories, from the point of ball release to the crossing of the vertical midline of the (x=0) position of the target (Figure 1B, first-order task). One of the trajectories corresponded exactly to the one they had induced with their movement, whereas the alternative one was determined by adding (or subtracting) from the velocity of release a given value (Δv), determined by an online staircasing procedure (see below). In a two-alternative forced-choice task (2AFC, first-order task), participants discriminated which of the two trajectories corresponded to the one that they had induced with their movement. To select a trajectory, they rotated the lever in either direction, and every 20° rotation would select a different trajectory (indicated on the screen by a thicker line). The order of which trajectory appeared thicker at the beginning of the first-order task was pseudo-randomised at the beginning of the experiment. Participants placed their index finger on the sensor to commit their response. This reporting led to long mean reaction times (RT mean ± SD: 2.23 ± 0.47 s). Immediately after the 2AFC decision, participants rated their confidence in their own discrimination response (second-order task) using a mouse to move a cursor on a continuous vertical scale ranging from very confident (top) to not confident (bottom). The starting position of the cursor on the continuous scale was pseudo-randomised per trial. Participants could indicate that they had made a procedural error (i.e., unintentionally selecting the wrong trajectory) by pressing the spacebar, and skipping the second-order task. They were instructed to report these errors only if they had made a procedural mistake and not when they were unsure of their answer. Those error trials (median (IQR: Q1-Q3): 2 (1-5) trials per participant) were excluded from the analyses.

#### 2.1.4 Experimental Manipulations

Each participant completed 240 trials that corresponded to one of two conditions: *congruent* or *incongruent*. The conditions differed on whether the alternative trajectory matched the real one in hitting or missing the target. More precisely, in the congruent condition, the alternative trajectory always had the same distal outcome as the real one. Simply put, if participants had hit (missed) the target with their ball throw, the alternative trajectory shown would have also hit (missed) the target. On the other hand, in the incongruent condition, the opposite was true: If participants had hit the target, the alternative ball trajectory shown would miss the target, and *vice versa*. Note that there is no linear mapping between trial congruency and velocity difference (Δv). Figure 1.D illustrates how the same Δv can lead to an alternative trajectory that is either congruent or incongruent with the actual ball trajectory. This resulted in a factorial 2 × 2 design with the factors of *Congruency* and *Outcome*. Participants first completed 16 training trials, followed by 16 feedback trials, that included trials for the two different conditions in pseudo-random order. On training trials, participants only threw the virtual ball and did the first-order, discrimination task, and — unlike during the experiment —, the target was visible throughout the ball flight. On feedback trials, participants additionally rated their confidence and received trial-wise feedback on their response accuracy in the first-order task: The cursor turned green or red following correct and incorrect responses, respectively.

#### 2.1.5 Online Staircases

The experiment included two (concurrent) online staircases. A 1-up, 1-down staircase adaptively determined the size of the ball and target, in order to keep participants’ rate of hitting the target at approximately 0.5. Additionally, a 2-down, 1-up staircase kept participants’ accuracy at approximately 71%, by controlling |Δv| (i.e., the absolute difference between the release velocity of the real and alternative trajectories). For any given trial, both the predefined condition (congruent/incongruent) and the ball throw outcome (target hit/miss) determined the alternative velocity. The alternative trajectory was computed by combining the |Δv| provided by the adaptive staircase with a predefined, pseudorandomised sign (+/-), resulting in the alternative trajectory appearing respectively to the right or left of the actual trajectory. If this did not lead to an alternative trajectory that matched the pre-defined condition, we deviated from the absolute Δv provided by the staircase as follows: first we changed the sign that would be combined with |Δv|. If this resulted in an alternative trajectory that did not match the pre-defined condition, we instead used the nearest absolute Δv value that met the condition. Note that this could result in stimuli presented more often to the left or to the right of the real trajectory or trials where the Δv was much smaller or much larger than the mean Δv provided by the staircase. We address these points in Experiment 2.

#### 2.1.6 Data Analysis

##### Exclusion Criteria

All data were excluded before any subsequent analysis steps. We excluded from the analysis trials where the reaction time (RT) for the first-order task was under 0.2 or above 8 seconds; trials where participants reported to have made a procedural error during the first-order task; trials that were trivially easy, meaning one of the trajectories hit the central pole and the other did not; and trials where the Δv used was not the Δv provided by the staircase and led to trials exceedingly easy or difficult compared to other trials of the same participant. During the analysis of the data, we implemented this pre-registered criterion by specifying as outliers the Δv that deviated more than 2 standard deviations (SDs) from the mean staircased value in the ten preceding trials. The median of trials excluded was 44 (IQR = 33-57) for each of the participants included in the final analysis. At pre-registration, we planned to exclude from the analysis the data from those participants that had response accuracy in the first-order task above 80% or below 60% within any given condition. Because this criterion would have led to excluding too many participants from the analyses, we decided to deviate from the pre-registered plan and make this threshold slightly more lenient (importantly, we made this decision before any statistical analyses on M-ratios). An upper bound of 85% accuracy is still considered reasonable to produce threshold performance and even higher values have been used elsewhere in metacognitive studies (e.g., Seow & Fleming, 2019). In our case this limit resulted in the exclusion of one participant. We also added one criterion to our pre-registered plan, and excluded from the analyses data from five participants due to a strong bias in the presentation of the stimuli: The real trajectory was presented to the right or left of the alternative on more than 70% of the trials. Datasets from six participants were excluded due technical issues that did not allow us to collect a full dataset, resulting in 34 participants being included in the analyses.

##### Estimates of Metacognitive Efficiency

To estimate metacognitive performance, we estimated metacognitive efficiency (M-ratio), which corresponds to metacognitive sensitivity (meta-d’) normalized by first-order sensitivity (d’) (Maniscalco & Lau, 2014). We first normalized confidence ratings by subtracting from the confidence values the minimal confidence rating of each condition and dividing by their range. We then discretized the normalized confidence ratings in 6 equidistant bins, adjusted for 0-count trials according to the default settings, and used the maximum likelihood estimation method in the MATLAB scripts provided on http://www.columbia.edu/∼bsm2105/type2sdt/. We then ran all statistical analyses in R (version 4.0.3, R Core Team, 2020). We used the BayesFactor package (Morey & Jeffrey, 2018) to obtain BF_10_ values.

### 2.2 Experiment 2

#### 2.2.1 Participants

For the follow-up Experiment 2, we recruited forty-two participants (27.8 ± 5.03 years old, 32 female) with the same inclusion and exclusion criteria (https://osf.io/javx5). Handedness data, collected post-hoc from thirty-five participants using the Edinburgh Handedness Inventory, confirmed that thirty-three participants were right-handed and two were left-handed (87 ± 26.2). Participants were all good English speakers, signed written informed consent before starting the study, and were compensated for their time with 8 €/hour. The study was approved by the ethics committee of the Institute of Psychology of the Humboldt-Universität zu Berlin and conducted according to the Declaration of Helsinki.

#### 2.2.2 Apparatus, Stimuli, and Procedure

The apparatus and stimuli were exactly as described for Experiment 1.

#### 2.2.3 Procedure

The procedure was as described for Experiment 1, save for the number of trials: Each participant completed 544 trials (split into four blocks) in the main experiment, as well as 40 training trials and eight feedback trials. Each experimental session took approximately two hours.

#### 2.2.4 Online Staircase

As in Experiment 1, we used a 1-up, 1-down staircase to adaptively determine the size of the ball and target. To better control the difficulty of the first-order task, in this follow-up experiment we used two separate 2-down, 1-up staircases to control the difference between the release and alternative velocity for the congruent and incongruent conditions. The alternative velocity was estimated for each condition in a similar fashion as in Experiment 1.

#### 2.2.5 Experimental Manipulations

The experimental manipulations were the same as in Experiment 1. In Experiment 2 we opted to adhere to the Δv provided by the staircase. In cases where the staircased Δv did not lead to the prespecified condition, we chose to maintain Δv and change the experimental condition instead. Note that this led to trial difficulties that were better staircased than in Experiment 1, but to an imbalance in the number of trials in each experimental condition (median, (IQR = Q1-Q2): congruent condition: 251, (233-259), incongruent: 225, (186-247) trials).

#### 2.2.6 Data Analysis

##### Exclusion Criteria

We followed the same exclusion criteria as in Experiment 1. We excluded 11 participants whose response accuracy in the first-order task was above 85 % or below 60% in one of the two conditions. We also excluded two participants due to a strong stimulus presentation bias. Finally, one participant could not complete all trials due to technical issues and their data were excluded from all analyses. The final sample size consisted of 28 participants The median of trials excluded was 67 (IQR = 40-110) trials from each participant.

##### Confirmatory analyses

We estimated M-ratios as described in Experiment 1. We ran parametric t-tests and two-way ANOVAS using the afex package (Singman et al, 2020) on normally distributed data. For the data that were not normally distributed, we ran Wilcoxon signed rank tests instead of t-tests and used the package ez for a non-parametric analysis of variance (ANOVA) (Lawrence, 2016).

##### Exploratory analyses

In addition to comparing first-order performance and metacognitive efficiency between conditions, we also studied metacognitive efficiency within each cell of the 2 × 2 factorial design. This resulted in relatively few trials per cell (median trial counts (IQR = Q1-Q3): congruent-hit = 145 (129-166), congruent-miss = 93 (78-117), incongruent-hit = 129 (100-164), incongruent-miss = 89 (66-98)). Because reliable estimates of meta-d’ using the MLE method have been shown to require at least 100 trials per condition (Fleming, 2017), we used the H-metad’ toolbox, a hierarchical Bayesian estimation of M-ratio estimation that is stable also for lower trial numbers (Fleming, 2017). We used default priors, three chains of 15.000 samples each, 5000 burn-in samples and a thinning parameter of 3. For all analyses, we visually inspected the chains for convergence and confirmed that the R-hat was approximately 1. We based our statistical inference on the degree of overlap between the 95% Highest Density Interval (HDI) of the difference between the posterior distributions on the one hand, and the region of practical equivalence (ROPE). We defined the limits of the ROPE as the interval around 0 with a half-width of 0.1 times the standard deviation of the pooled M-ratios from the confirmatory analysis (Kruschke, 2018).

## 3. Results

To test whether the outcome of a movement informs motor metacognitive judgments, we compared metacognitive efficiency between two conditions that differed on the type of information available for the discrimination task: In the congruent condition both trajectories had the same distal outcome (hit/miss the target). Therefore, the only information available to make the discrimination decision was the entire ball trajectory as represented on the screen, that resulted from different ball release velocities. In the incongruent condition, the trajectories had different distal outcomes: When participants hit (missed) the target the alternative trajectory missed (hit) it. This meant that on incongruent trials participants had additional information about the distal outcome of the action, and could therefore use this additional piece of information to make their decisions.

### 3.1 Experiment 1

#### 3.1.1. Confirmatory analyses

We evaluated separately zero-order (motor), first-order, and second-order performance. Participants performed well in the zero-order motor task: Despite the 1-up, 1-down staircase, aimed at achieving a rate of target hit of 0.5, participants more often hit than missed the target (mean hit rate: 0.61 ± 0.10). We found weak evidence for improvement in motor performance over time (p = 0.002, BF_10_ = 0.65; See SI for details). An analysis of performance in the first-order task confirmed that the experimental manipulation had the expected effect: Participants were better able to discriminate the actual from the alternative trajectories in the incongruent (d’ = 1.67 ± 0.19) compared to the congruent condition (d’ = 1.02 ± 0.2; t(33) = -13.42, p < 0.001, Cohen’s d = -2.3, BF_10_ = 1.02 ×10^12^, Figure 2.A). An analysis of reaction times revealed a similar pattern, with shorter RTs in the incongruent condition (RT = 2.17 ± 0.45 s) as compared to the congruent condition (RT = 2.3 ± 0.5; for RTs transformed (1/x): t(33) = - 2.98, p < 0.01, Cohen’s d = -0.511, BF_10_ = 7.35). This suggests that, as expected by experimental design, participants used information on the outcome to solve the discrimination task, which was only possible on incongruent trials.

**Figure 2:**
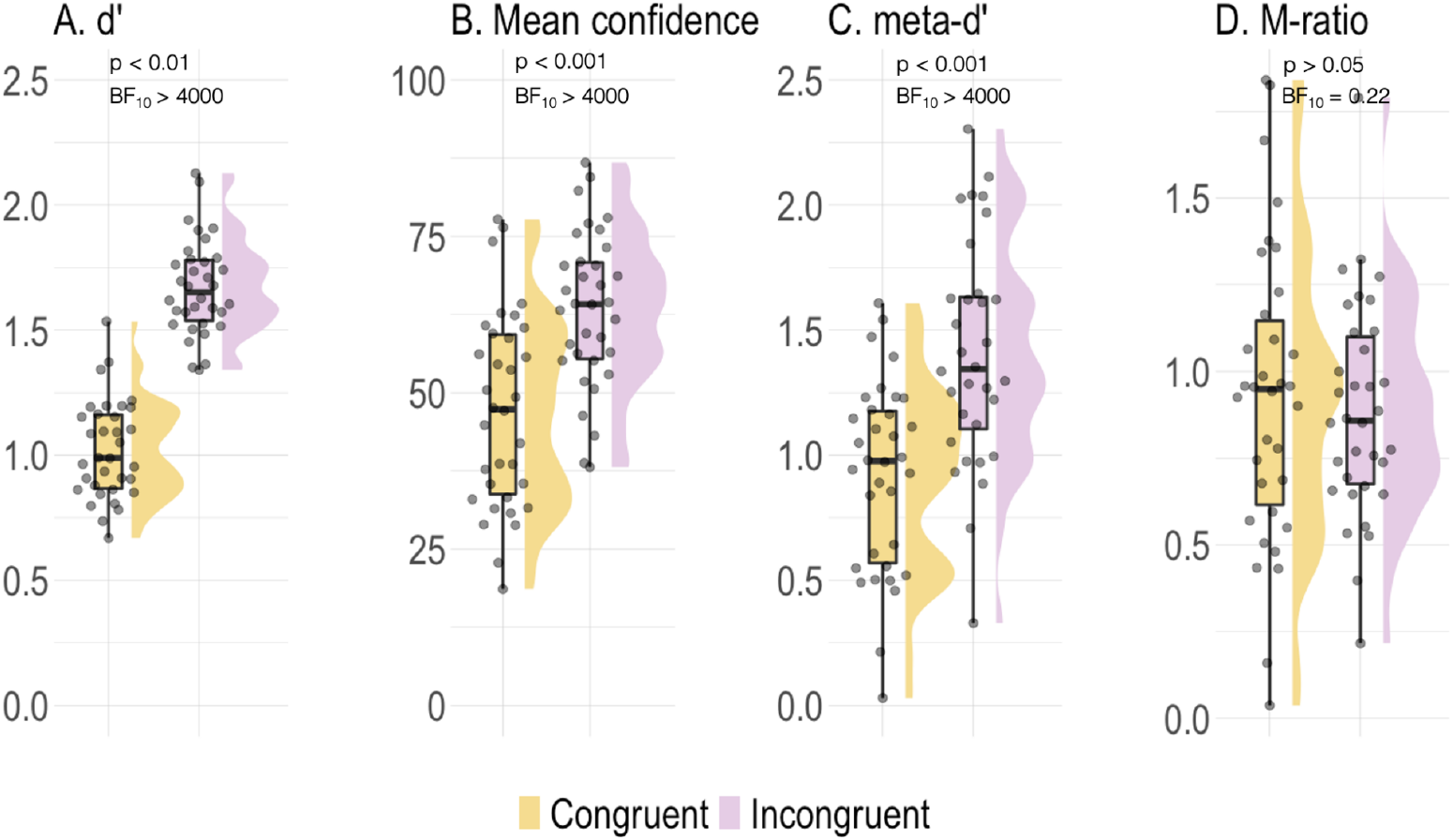
First- and second-order performance measures for Experiment 1. The violin plots depict the smoothed distribution of the data for four main summary measures: **(A.) d’:** first-order performance in the first-order task, **(B.) Mean Confidence, (C.) meta-d’:** Metacognitive sensitivity and **(D.) M-ratio:** Metacognitive efficiency. Each dot represents a single participant. The overlaid box plots represent the interquartile range. d’, Confidence ratings and meta-d’ were significantly higher for the incongruent condition. We found no differences in metacognitive efficiency (M-ratio) between congruent and incongruent conditions.

In line with higher first-order performance, mean confidence ratings were also higher in the incongruent condition (63.35 ± 12.37) compared to the congruent condition (47.15 ± 15.54; t(33) = -10.149, p < 0.001, Cohen’s d = -1.74, BF_10_ = 8.14 ×10^10^, Figure 2.B). Crucially, an analysis of second-order performance measures revealed that, while participants’ metacognitive sensitivity (meta-d’) was higher (t(33) = -6.4026, p < 0.001; Cohen’s d = -1.10, BF_10_ = 5.4 ×10^4^, Figure 2C) in the incongruent condition (1.48 ± 0.53) compared to the congruent condition (0.92 ± 0.38), M-ratio (which, unlike metad’, controls for first-order performance) did not differ between conditions (M-ratio incongruent = 0.89 ± 0.31; M-ratio congruent = 0.93 ± 0.43; t(33) = 0.651, p > 0.05, Cohen’s d = 0.112, BF_10_ = 0.22, Figure 2.D). Together, these results suggest that the outcome information, while being beneficial for the first-order task, did not provide any additional advantage for metacognitive judgments.

While the absence of differences in metacognitive efficiency is interpretable because M-ratio controls for first-order performance, the large difference in first-order d’ may still be problematic and it is preferable to compare conditions with equal or more similar first-order performance. Moreover, we noted that the difficulty of the discrimination decision differed between congruency conditions (congruent condition: Δv = 0.17 ± 0.08, incongruent condition: Δv = 0.27 ± 0.13; Δv 1/x transformed: t(33) = 8.83, p < 0.001, Cohen’s d = 1.51, BF_10_ = 3.2 ×10^7^). Hence, in Experiment 1 neither stimulus presentation, in terms of the trial difficulty, nor first-order performance were optimally controlled. We conducted the follow-up Experiment 2, addressing this confound by using two separate staircases for the two conditions to ensure better experimental control.

### 3.2 Experiment 2

#### 3.2.1. Confirmatory Analyses

As in Experiment 1, we found that participants hit the target in the majority of the trials (target hit rate = 0.68 ± 0.15), despite the online staircases to control the ball and target size. Unlike in experiment 1, we found no significant changes in motor performance over time, only a small but non-significant decrease in the distance to the target as trials progressed (See SI for details). A paired samples t-test revealed that Δv were higher on incongruent trials (0.47 ± 0.24 m/s) compared to congruent trials (0.41 ± 0.21; Δv (1/x transformed): t(27) = 5.04, p < 0.01, Cohen’s d = 0.95, BF_10_ = 859). This difference in Δv between conditions was in the same direction, but with a smaller effect size, than what we found in Experiment 1.

##### Effect of Congruency

In the discrimination task, participants were more accurate on incongruent trials (d*’ =* 1.68 ± 0.44, Figure 3.A) compared to congruent trials (d*’ =* 1.46 ±0.2: Wilcoxon Signed-Ranks test: Z = -3.03, p < 0.05, r = 0.87), despite task difficulty now being controlled by two separate staircases for these conditions. Participants were also faster to respond on incongruent (RT = 1.94 ± 0.37 s) compared to congruent trials (RT = 2 ± 0.39 s: Wilcoxon Signed-Ranks Test: Z = 2.69, p < 0.01, r = 0.36). Accordingly, mean confidence ratings were higher on incongruent trials (71.29 ± 14.03) compared to congruent trials (68.63 ±15.08; t(27) = -4.24, p < 0.001, Cohen’s d = -0.8, BF_10_ = 121.2, Figure 3.B). As in Experiment 1, meta-d’ was also higher in the incongruent condition (1.22 ± 0.48) compared to the congruent condition (0.9 ± 0.5; t(27) = -4.3, p < 0.01, Cohen’s d = -0.82, BF_10_ = 151.1, Figure 3.C) but, in line with the results from Experiment 1 and against our hypothesis, we found no evidence for a difference in the M-ratio values (M-ratio incongruent = 0.74 ± 0.28; M-ratio congruent = 0.62 ± 0.38; t(27) = -1.79, p > 0.05, Cohen’s d = -0.34, BF_10_ = 0.82, Figure 3.D).

**Figure 3:**
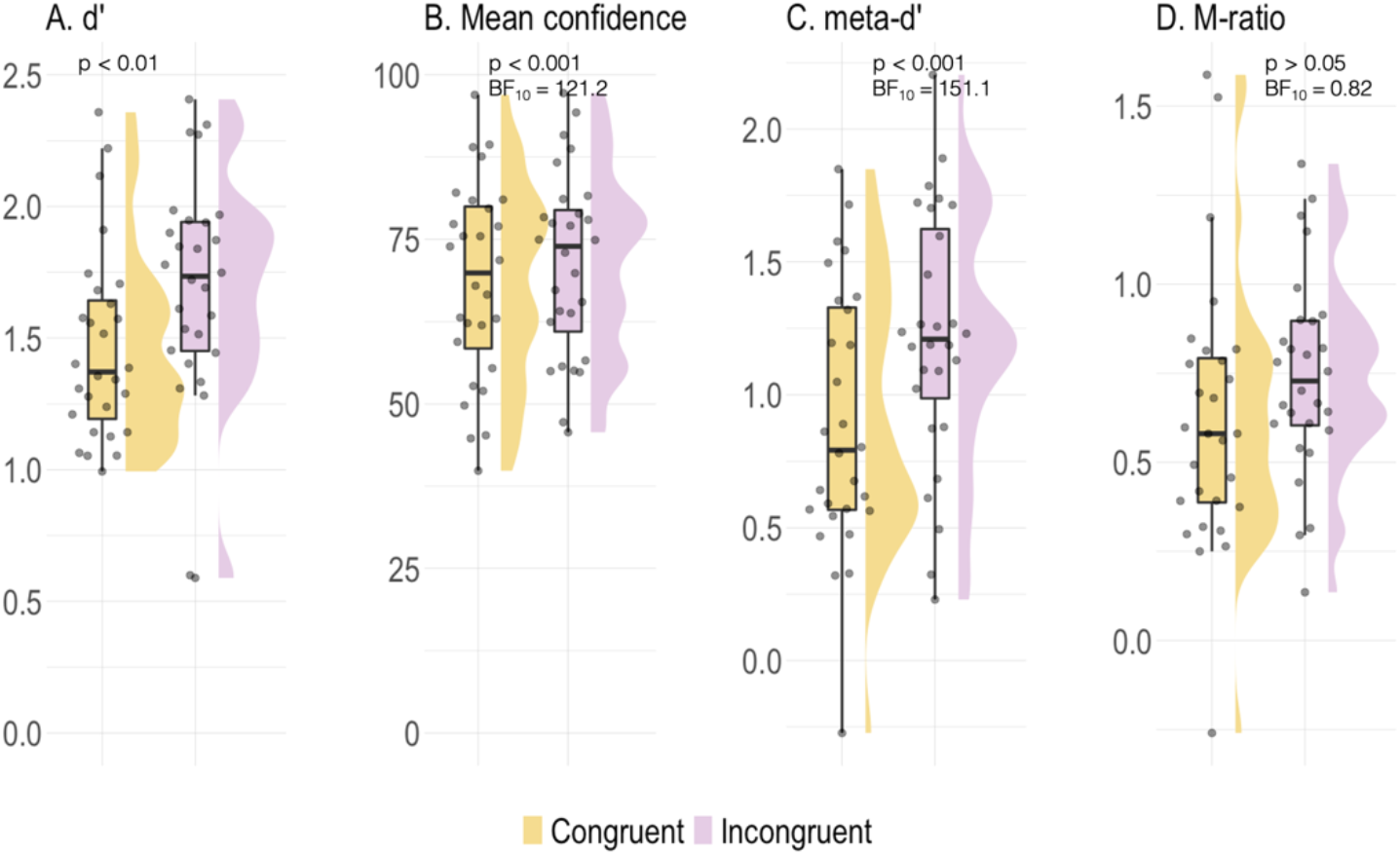
First- and second-order performance measures for Experiment 2. The violin plots illustrate the smoothed distributions of the data for four main summary measures: **(A.) d’:** first-order performance in the first-order task, **(B.) Mean Confidence, (C.) meta-d:** Metacognitive sensitivity, and **(D.) M-ratio** Metacognitive efficiency. Each dot represents estimates for a single participant for any given condition. The overlaid box plot indicates the interquartile range. d’ measures performance in the discrimination task. d’, Mean confidence ratings and meta-d’ were significantly higher for the incongruent condition. As in Experiment 1, we found no differences in metacognitive efficiency (M-ratio) between conditions.

#### 3.2.2 Exploratory analyses

##### Interactions between Outcome and Congruency on metacognitive efficiency

All confirmatory analyses focused on the effects of *Congruency*, but collapsed across *Outcome*. In this set of exploratory analyses, we address the impact of *Outcome*, namely hitting the target on first- and second-order responses, and its interactions with *Congruency*. We conducted these analyses exclusively on the data from Experiment 2 because it included more trials, and better controlled performance, as compared to Experiment 1.

##### Effects of Outcome and Congruency on first-order performance

We first investigated performance in the first-order task. A non-parametric ANOVA on d’ revealed a main effect of *Congruency* (p < 0.05) and a main effect of *Outcome* (p < 0.05, Figure 4.A). Participants performed better on the first-order task on incongruent trials (d’ = 1.72 ± 0.92) than congruent trials (d’ = 1.41 ± 0.49). Additionally, participants performed better when they hit the target (d’ = 1.83 ± 0.62) compared to when they missed it (d’ = 1.30 ± 0.78). Interestingly, participants’ first-order performance varied greatly in the incongruent compared to the congruent condition in trials in which the ball missed the target. In fact, some participants’ d’ values were even close to or below zero (Figure 4.A). A possible explanation for these cases is that, on those trials where the ball missed the target, participants showed a hit bias: They disregarded their actual motor performance and selected a trajectory that implied a target hit in line with their intention. The results reported here include the data from the four participants with d’ below zero, but excluding them led to the same pattern of results (see Supplementary material).

**Figure 4:**
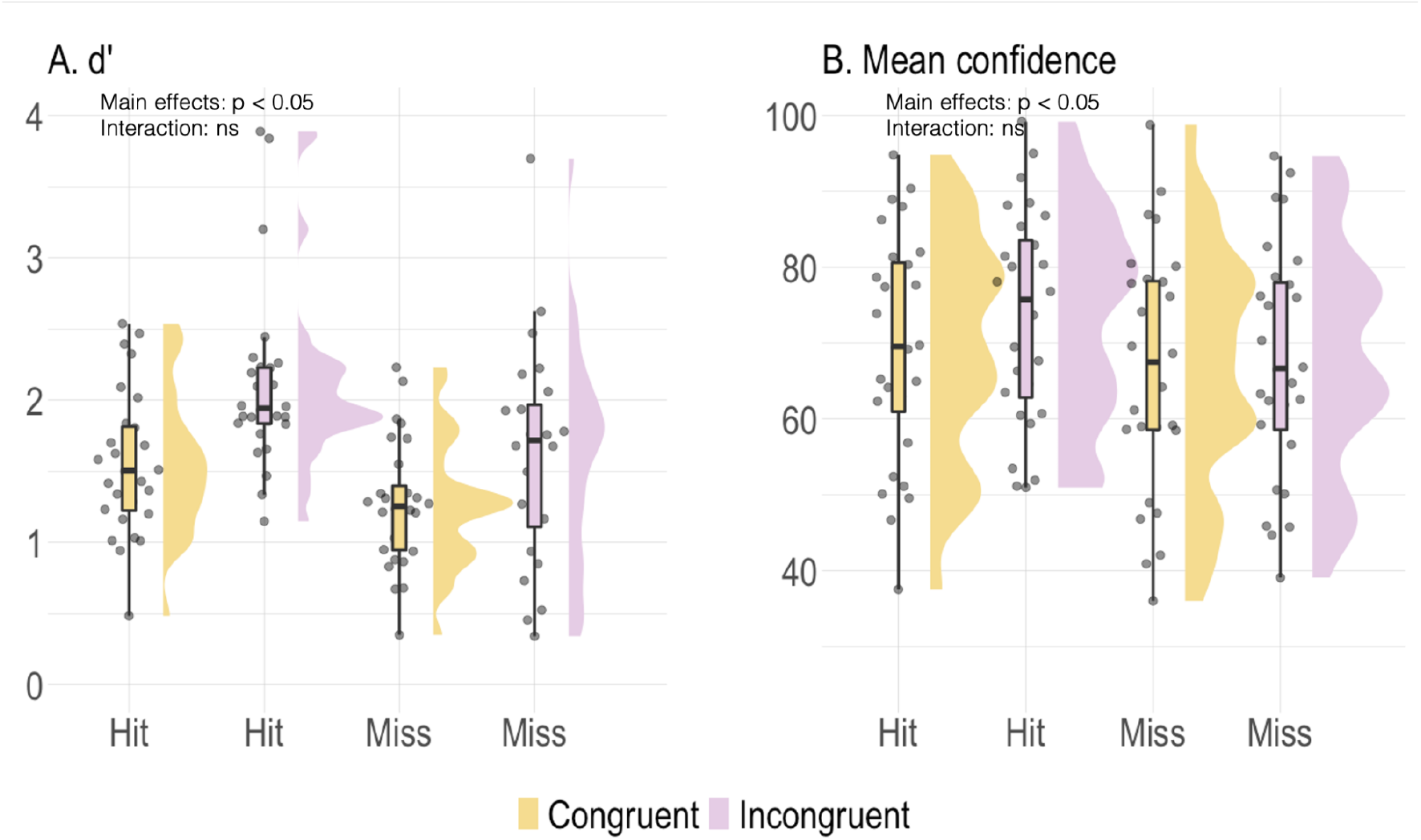
Effects of *Congruency* and *Outcome* on first-order performance and confidence ratings. The violin plots illustrate the smoothed distributions of the data split according to outcome (hit/miss the target) and condition (congruent/incongruent). Each dot represents a single participant. The overlaid box plot indicates the interquartile range. d’ measures performance in the discrimination task. Main effects are marked next to the plots with an asterisk. None of the interaction effects were significant.

##### Effect of Outcome and Congruency on confidence

A two-way ANOVA on mean confidence revealed a main effect of *Outcome* (F(1,27) = 18.82, p < 0.001, BF_10_ = 656) and a main effect of *Congruency* (F(1,27) = 13.93, p < 0.001, BF_10_ = 2.43) but no interaction (F(1,27) = 1.7, p = 0.2, BF_10_ = 0.69). These results mirror those of first-order performance: Participants were simply generally more confident in those conditions where they were more often correct.

##### Effect of Outcome and Congruency on second-order performance

We then examined potential interactions between the effects of *Outcome* and *Congruency* on metacognitive efficiency. Because each cell of the factorial 2 × 2 design included relatively few trials (median trial counts (IQR = Q1-Q3): congruent-hit = 145 (129-166), congruent-miss = 93 (78-117), incongruent-hit = 129 (100-164), incongruent-miss = 89 (66-98)), we estimated M-ratios using the HMetad’ toolbox (Fleming, 2017). A two-way ANOVA revealed that the 95% HDI [-0.51, 0.07] of the interaction effect (Outcome × Congruency) slightly overlapped with zero (Figure 5.A). Therefore, we examined the differences between pairs to further understand this result. Pairwise comparisons revealed that, for those trials where participants hit the target, M-ratio estimates were higher for incongruent trials than for congruent trials: The 95% HDI of the difference [0.58, 0.08] (incongruent minus congruent) excluded the ROPE [-0.034, 0.034] (Figure 5.B). This was not the case for trials where participants missed the target, where there was no advantage of *Congruency* on M-ratios: the 95% HDI of the difference [-0.36, 0.5] overlapped with zero and the ROPE [-0.034, 0.034] (Figure 5.C). These results indicate that the outcome information is beneficial specifically for metacognitive judgments, even after controlling for first-order performance, only when participants reach the motor goal of hitting the target, but not when they missed it. We wondered whether attentional effects could offer a parsimonious explanation for this pattern of results: Participants could have been more likely to both hit the target and provide more precise confidence ratings on those trials where they were more attentive. Crucially, if this were the case, we would also expect shorter RTs on these trials. However, the data do not support this explanation, neither on first-or second-order responses. A non-parametric ANOVA on first-order RTs revealed a main effect of condition (p < 0.05), driven by faster discrimination responses on incongruent trials (1.94 ± 0.38 s) compared to congruent trials (2.01 ± 0.43 s), but no main effect of *Outcome* (p > 0.05) or interaction with *Congruency* (p > 0.05). Similarly, a two-way ANOVA on the (1/x transformed) reaction times of the second-order task, revealed a main effect of *Congruency* (F(1,27) = 11.87, p < 0.01, BF_10_ = 68.76 × 10^3^), no main effect of *Outcome* (F(1,27) = 0.99, p = 0.33, BF_10_ =0.43) and no interaction (F(1,27) = 0.55, p = 0.46, BF_10_ = 0.31). This indicates the *Congruency* advantage when participants hit the target cannot be simply explained by attentional effects.

**Figure 5:**
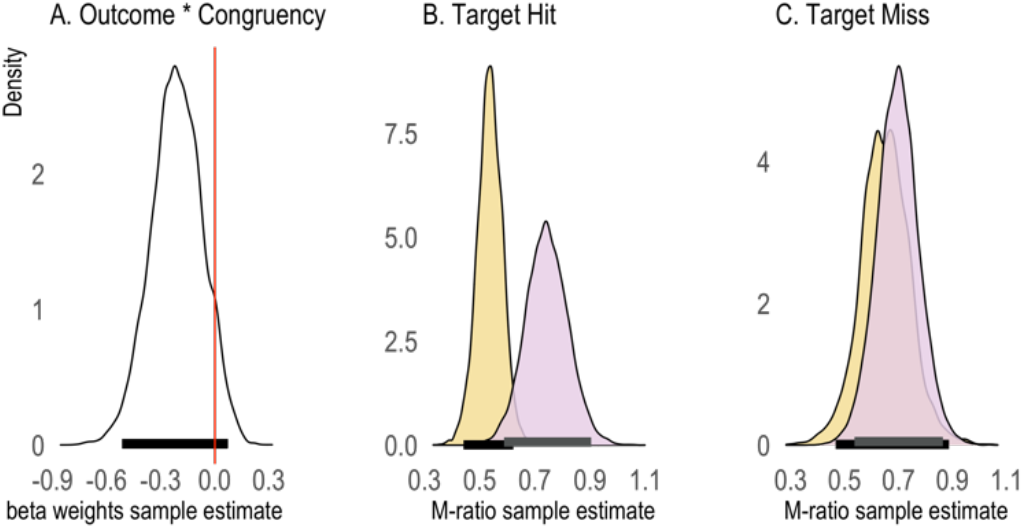
Effects of *Congruency* and *Outcome* on metacognitive efficiency (n = 28). **(A.)** Group posterior samples from the beta value coding the interaction *Outcome* × *Congruency*. **(B.)** Group posterior estimates of metacognitive efficiency (M-ratio) for congruent (yellow) and incongruent (pink) conditions for those trials where participants hit the target. **(C.)** Group posterior estimates of metacognitive efficiency (M-ratio) for congruent (yellow) and incongruent (pink) conditions for those trials where participants missed the target. The black and gray lines indicate the highest density interval (HDI) and the dashed line, the region of practical equivalence (ROPE).

## Discussion

In two experiments, we asked whether the distal outcome of a goal-oriented movement informs metacognitive representations of that movement. Following a now widespread operationalization (Locke et al., 2020), we quantified metacognitive performance as the relationship between confidence ratings and accuracy in a discrimination task. With a quick arm-movement, participants threw a virtual ball, then chose which of two trajectories best corresponded to their movement, and rated their confidence in their preceding binary choice. We included two conditions that differed on whether the distal outcome of the movement was informative or not for the discrimination decision. In the *congruent* condition, the two trajectories led to the same distal outcome (both hit or both missed the target), therefore the outcome was not informative. In the *incongruent* condition, the two trajectories differed not only on low-level parameters, but also led to different distal outcomes (one hit and the other missed the target). Hence, movement outcome was an additional piece of information, available on incongruent, but not congruent trials.

We found that population mean M-ratios were above zero in both conditions. This suggests that participants could metacognitively access their own throwing movements and is in line with previous research showing above-chance metacognitive ability in describing voluntary movements (Arbuzova et al., 2021; Charles et al., 2020; Locke et al., 2020; Mole et al., 2018; Pereira et al., 2021; Sinanaj et al., 2015).

### Effects of congruency information

Following our pre-registered plan, we first compared congruent and incongruent conditions, regardless of the actual outcome (i.e., we pooled together trials where participants had hit or missed the target with their throw). Across the two experiments, we consistently found that participants performed better in the first-order discrimination task on incongruent trials, where the two alternative trajectories represented two ball flights that led to a different outcome, as compared to congruent trials. This suggests that information about the outcome sharpened first-order representations and is in agreement with a previous study (David et al, 2016) showing that participants’ agency ratings (which we argue to be equivalent to first-order representations, see below) were more sensitive to feedback delays applied to the outcome of a movement than to a representation of the movement. However, and surprisingly, we did not find evidence that outcome information in general improved metacognitive efficiency in either one of the two experiments. This result goes against our pre-registered hypothesis and suggests that outcome information is not the primary factor that informs metacognitive motor representations. Instead, metacognitive representations might rely on both the low-level motor parameters and distal outcome as sources of partially redundant information. This recalls previous studies where no differences in second-order precision were evident despite clear differences in the kind of information available for the first-order task. Neither active vs. passive movements (Charles et al., 2020) nor the monitoring of the amplitude vs. the speed of a movement (Arbuzova et al., 2021) yielded differences in metacognitive efficiency.

### Interactions between the effects of Outcome and Congruency

In exploratory analyses, we tested how outcome information affects metacognitive representations on successful and unsuccessful trials. We carried out this analysis only on the data from the second experiment, in which we used condition-specific adaptive procedures to control participants’ performance and where more trials were available for each condition. When we quantified metacognitive efficiency separately for trials in which the ball hit or missed the target, we found an interesting pattern of results which, we note, should be interpreted with caution given the generally low number of trials available in which participants missed the target. A low number of trials can lead to unreliable estimates of meta-d’, which we aimed to prevent by using the Hmetad’ toolbox (Fleming, 2017). For first-order performance, we found that participants were better able to discriminate between the two trajectories when they had hit the target, as compared to when they had missed it. In other words, we found no interaction between Congruency and Outcome at the first-order level: The effect of congruency on d’ that we discussed in the previous section, was not modulated by the distal outcome. This is in line with previous literature suggesting that the information of whether a movement reached a goal affects motor representations (Blakemore et al., 2002; Fourneret & Jeannerod, 1998; Gaveau et al., 2014). It also aligns with previous studies of the effect of outcome manipulations on the human sense of agency. For instance, in a previous study, participants moved their fingers to touch one of two possible targets and judged the synchronicity between their own finger movements and a virtual hand.

These judgments indirectly measure participants’ sense of agency (Villa et al., 2018), and were affected by experimental manipulations of both delay in the movement and touching the correct target, suggesting that aside from motor parameters, the information of whether a movement reached a goal does affect the subjective experience of control.

Beyond the well-established effects of outcome on motor representations, our paradigm allowed us to decouple motor intentions from reaching a goal. Unlike in simple finger-movement tasks, participants were not always successful in hitting the target despite presumably always intending to. Thus, these results suggest that the contribution of goal-related information to higher-order motor representations might interact with intentions. In our experiment, incorrect discrimination responses on incongruent trials when the ball missed the target amount to participants misattributing a successful movement to themselves, and (falsely) reporting that the movement they just made led to the virtual ball hitting the target, when in fact it had not. Intriguingly, we found evidence for a hit bias, that is participants often made these misattribution errors. Indeed, in this condition only, some participants even had negative d’ values (i.e., their performance was below chance) on incongruent trials when they missed the target. This behavior is compatible with the theory of mental causation, according to which the agent infers that they control an action based on whether the action matched their intentions (Wegner & Wheatley, 1999) and is also compatible with more recent results showing that experienced typists failed to explicitly report committed mistakes that were automatically corrected by the experimenters (unbeknownst to the participants), because the output on the screen matched their intention (Logan & Crump, 2010). Therefore, we speculate that participants relied on predictions based on their motor intentions to decide which trajectory represented their own.

At the metacognitive level we found that outcome information was advantageous only when participants had successfully hit the target. That is, on hit but not on miss trials, metacognitive representations were more precise on *incongruent* as compared to *congruent* trials, above and beyond what would be expected given differences in first-order performance. To interpret this result, let us first assume that participants’ intention on each trial was to hit the target (in keeping with the task instructions) and that they consequently formed an expectation that the ball’s trajectory would hit the target. Then, it follows that on incongruent trials participants discriminated between an expected trajectory that matched their proximal intentions (Mylopoulos & Pacherie, 2019), regardless of their *actual* motor performance, and an unexpected one. On hit trials, congruent and incongruent conditions differed on whether participants discriminated between two trajectories that both had expected outcomes (both hit) or one that did, and another that did not. On the other hand, on miss trials, congruent and incongruent conditions differed in that either both trajectories (congruent condition) or only the incorrect alternative (incongruent condition) were unexpected. Note that this interpretation relies on participants being able to separately monitor motor intentions and motor execution, which has been recently shown in a time estimation task (Frömer et al., 2021). The increase in metacognitive efficiency that we observed, specific to incongruent hit trials (where the movement outcome matched the motor intention), links prior expectations and metacognitive efficiency. At least two previous studies showed complementary effects of expectations on the precision of metacognitive representations. On the one hand, prior expectations have been shown to enhance metacognitive representations when the decision is congruent with expectations in a perceptual decision task (Sherman et al., 2015). On the other hand, unexpected perceptual outcomes of an action have been argued to lead to enhanced metacognition (Yon, 2020). Our results contribute to this growing literature examining the interactions between prior expectations and metacognitive representations.

### Future directions: Methods to study the sense of agency

The findings from this study relate to research on the sense of agency, namely the subjective experience that we are the authors of our movements and actions (Haggard, 2017; Moore, 2016). ‘Agency’ is used as an umbrella term and is neither conceptually nor operationally strictly defined (Charalampaki et al., 2022; David, 2012; Gallagher, 2012; Haggard, 2017; Pacherie, 2007; Synofzik et al., 2008). To study the human sense of agency, researchers often ask participants to make a movement, and display the movement and their consequences back to participants. Experimental manipulations then alter either the representation of the movement itself or the movement outcome, who then explicitly rate their subjective experience (Grünbaum & Christensen, 2020). In these paradigms, participants are effectively asked to detect trials in which they experience the loss of agency, and respond “Yes” or “No” to the question of whether they felt they were the agents of movement. Here, we replaced a detection task with a discrimination task that is computationally equivalent. Indeed, we have discussed the effects of congruency that we observed on first-order performance in light of previous studies on the sense of agency (Villa, 2018; David, 2016), in keeping with recent findings suggesting that judgments of agency reflect first- and not second-order processing (Constant et al., 2021). A future interesting research direction may be to study the sense of agency using a similar experimental protocol, that would allow for direct comparisons of sensitivity between agency following perturbations of different levels of a movement (proximal vs. distal outcome, body vs. corresponding movement, Dogge et al., 2019).

## Conclusion

We found that young, healthy participants can accurately metacognitively monitor their motor performance. Further, both first-order performance and metacognitive efficiency were higher when the outcome of an action matched participants’ intentions, as compared to when these did not match. On the basis of these results, we suggest a central role for motor intentions in metacognitive motor representations with an over-reliance on motor intentions in detriment of motor monitoring.

## Author contributions

**Angeliki Charalampaki:** Conceptualization, Data curation, Formal analysis, Investigation, Methodology, Project administration, Software, Validation, Visualization, Writing – original draft, Writing -review & editing. **Caroline Peters:** Conceptualization, Data curation, Investigation, Methodology, Project administration, Software, Writing - review & editing. **Heiko Maurer:** Methodology, Software, Writing - review & editing. **Lisa K. Maurer:** Methodology, Writing - review & editing. **Hermann Müller:** Methodology, Writing - review & editing. **Julius Verrel:** Conceptualization, Writing - review & editing. **Elisa Filevich:** Conceptualization, Formal analysis, Funding acquisition, Methodology, Project administration, Resources, Software, Supervision, Validation, Writing – original draft, Writing - review & editing.

## Acknowledgements

AC was supported by the Deutscher Akademischer Austauschdienst (DAAD). AC, CP, and EF were supported by a Freigeist Fellowship to EF from the Volkswagen Foundation (grant number 91620). LKM and HM were supported by DFG, German Research Foundation (project 222641018 - SFB/TTR 135 TP B6). The funders had no role in the conceptualization, design, data collection, analysis, decision to publish, or preparation of the manuscript. We would like to thank Anthony Ciston and Christina Koß for help with data collection.

## Web references

The scripts required for estimating the M-ratio using the MLE method were downloaded from http://www.columbia.edu/∼bsm2105/type2sdt/ (as accessed September 2020).

All R scripts for the hierarchical Bayesian estimation of M-ratio (meta-d’/d’) were downloaded from https://github.com/metacoglab/HMeta-d (as accessed April 2022).

## Code availability

The raw data, as well as MATLAB and R scripts to reproduce the analysis and figures are freely available at https://gitlab.com/AngelikiC/metacognition-of-outcome-with-skittles.

## Bibliography

Arbuzova, P., Peters, C., Röd, L., Koß, C., Maurer, H., Maurer, L. K., Müller, H., Verrel, J., & Filevich, E. (2021). Measuring metacognition of direct and indirect parameters of voluntary movement. Journal of Experimental Psychology: General. https://doi.org/10.1037/xge0000892

Blakemore, S.-J., Wolpert, D. M., & Frith, C. D. (2002). Abnormalities in the awareness of action. Trends in Cognitive Sciences, 6(6), 237–242. https://doi.org/10.1016/S1364-6613(02)01907-1

Bourdin, P., Martini, M., & Sanchez-Vives, M. V. (2019). Altered visual feedback from an embodied avatar unconsciously influences movement amplitude and muscle activity. Scientific Reports, 9(1), 19747. https://doi.org/10.1038/s41598-019-56034-5

Brainard, D. H. (1997). The Psychophysics Toolbox. Spatial Vision, 10(4), 433–436. https://doi.org/10.1163/156856897X00357

Charalampaki, A., Karabanov, A.N., Ritterband-Rosenbaum, A., Nielsen, J.B., Siebner, H.R., Christensen, M.S. (2022). Sense of agency as synecdoche: Multiple neurobiological mechanisms may underlie the phenomenon summarized as sense of agency. Consciousness and Cognition (In Press)

Charles, L., Chardin, C., & Haggard, P. (2020). Evidence for metacognitive bias in perception of voluntary action. Cognition, 194, 104041. https://doi.org/10.1016/j.cognition.2019.104041

Constant, M., Salomon, R., & Filevich, E. (2022). Judgments of agency are affected by sensory noise without recruiting metacognitive processing. Elife, 11, e72356.

David, N., Skoruppa, S., Gulberti, A., Schultz, J., & Engel, A. K. (2016). The sense of agency is more sensitive to manipulations of outcome than movement-related feedback irrespective of sensory modality. PLoS One, 11(8), e0161156.

David, N. (2012). New frontiers in the neuroscience of the sense of agency. Frontiers in Human Neuroscience, 6, 161. https://doi.org/10.3389/fnhum.2012.00161

Dogge, M., Custers, R., & Aarts, H. (2019). Moving Forward: On the Limits of Motor-Based Forward Models. Trends in Cognitive Sciences, 23(9), 743–753. https://doi.org/10.1016/j.tics.2019.06.008

Fleming, S. M. (2017). HMeta-d: Hierarchical Bayesian estimation of metacognitive efficiency from confidence ratings. Neuroscience of Consciousness, 2017(nix007). https://doi.org/10.1093/nc/nix007

Fleming, S. M., & Lau, H. C. (2014). How to measure metacognition. Frontiers in Human Neuroscience, 8, 443. https://doi.org/10.3389/fnhum.2014.00443

Fourneret, P., & Jeannerod, M. (1998). Limited conscious monitoring of motor performance in normal subjects. Neuropsychologia, 36(11), 1133–1140. https://doi.org/10.1016/S0028-3932(98)00006-2

Frömer, R., Nassar, M. R., Bruckner, R., Stürmer, B., Sommer, W., & Yeung, N. (2021). Response-based outcome predictions and confidence regulate feedback processing and learning. ELife, 10, e62825. https://doi.org/10.7554/eLife.62825

Gallagher, S. (2012). Multiple aspects in the sense of agency1. New Ideas in Psychology, 30(1), 15–31. https://doi.org/10.1016/j.newideapsych.2010.03.003

Gaveau, V., Pisella, L., Priot, A.-E., Fukui, T., Rossetti, Y., Pélisson, D., & Prablanc, C. (2014). Automatic online control of motor adjustments in reaching and grasping. Neuropsychologia, 55, 25–40. https://doi.org/10.1016/j.neuropsychologia.2013.12.005

Grünbaum, T., & Christensen, M. S. (2020). Measures of agency. Neuroscience of Consciousness, 2020(1). https://doi.org/10.1093/nc/niaa019

Haggard, P. (2017). Sense of agency in the human brain. Nature Reviews Neuroscience, 18(4), 196–207. https://doi.org/10.1038/nrn.2017.14

Joch, M., Hegele, M., Maurer, H., Müller, H., & Maurer, L. K. (2017). Brain negativity as an indicator of predictive error processing: The contribution of visual action effect monitoring. Journal of Neurophysiology, 118(1), 486–495. https://doi.org/10.1152/jn.00036.2017

Kleiner, M., Brainard, D., Pelli, D., Ingling, A., Murray, R., & Broussard, C. (2007). What’s new in Psychtoolbox-3. Perception, 36(14), 1–1.

Kruschke, J. K. (2018). Rejecting or Accepting Parameter Values in Bayesian Estimation. Advances in Methods and Practices in Psychological Science, 1(2), 270–280. https://doi.org/10.1177/2515245918771304

Latash, M. (2000). There is no motor redundancy in human movements. There is motor abundance. Motor Control, 4(3), 259–260.

Latash, M. L. (2012). The Bliss of Motor Abundance. Experimental Brain Research. Experimentelle Hirnforschung. Experimentation Cerebrale, 217(1), 1–5. https://doi.org/10.1007/s00221-012-3000-4

Lawrence, M. A. (2016). ez: Easy Analysis and Visualization of Factorial Experiments. R package version 4.4-0. https://CRAN.R-project.org/package=ez

Locke, S. M., Mamassian, P., & Landy, M. S. (2020). Performance monitoring for sensorimotor confidence: A visuomotor tracking study. Cognition, 205, 104396. https://doi.org/10.1016/j.cognition.2020.104396

Logan, G. D., & Crump, M. J. C. (2010). Cognitive Illusions of Authorship Reveal Hierarchical Error Detection in Skilled Typists. Science. https://www.science.org/doi/abs/10.1126/science.1190483

Makowski, D., Ben-Shachar, M., & Lüdecke, D. (2019). bayestestR: Describing Effects and their Uncertainty, Existence and Significance within the Bayesian Framework. Journal of Open Source Software, 4(40), 1541. doi:10.21105/joss.01541

Maniscalco, B., & Lau, H. (2014). Signal Detection Theory Analysis of Type 1 and Type 2 Data: Meta-d′, Response-Specific Meta-d′, and the Unequal Variance SDT Model. In S. M. Fleming & C. D. Frith (Eds.), The Cognitive Neuroscience of Metacognition (pp. 25–66). Springer. https://doi.org/10.1007/978-3-642-45190-4_3

Maurer, L. K., Maurer, H., & Müller, H. (2015). Neural correlates of error prediction in a complex motor task. Frontiers in Behavioral Neuroscience, 9, 209. https://doi.org/10.3389/fnbeh.2015.00209

Maurer, L.K., Maurer, H., Hegele, M., Müller, H. (2022) Can Stephen Curry really know?—Conscious access to outcome prediction of motor actions. PLoS ONE 17(1): e0250047. https://doi.org/10.1371/journal.pone.0250047

Miall, R. C., & Wolpert, D. M. (1996). Forward Models for Physiological Motor Control. Neural Networks, 9(8), 1265–1279. https://doi.org/10.1016/S0893-6080(96)00035-4

Mole, C. D., Jersakova, R., Kountouriotis, G. K., Moulin, C. J., & Wilkie, R. M. (2018). Metacognitive judgements of perceptual-motor steering performance. Quarterly Journal of Experimental Psychology, 71(10), 2223–2234. https://doi.org/10.1177/1747021817737496

Moore, J. W. (2016). What Is the Sense of Agency and Why Does it Matter? Frontiers in Psychology, 7, 1272. https://doi.org/10.3389/fpsyg.2016.01272

Morey, R. D., & Rouder, J. N. (2018). BayseFactor: computation of bayes factors for common designs. R package version 0.9.12-4.2. https://CRAN.R-project.org/package=BayesFactor

Müller, H., & Sternad, D. (2004). Decomposition of Variability in the Execution of Goal-Oriented Tasks: Three Components of Skill Improvement. Journal of Experimental Psychology: Human Perception and Performance, 30(1), 212–233. https://doi.org/10.1037/0096-1523.30.1.212

Mylopoulos, M., & Pacherie, E. (2019). Intentions: The dynamic hierarchical model revisited. WIREs Cognitive Science, 10(2), e1481. https://doi.org/10.1002/wcs.1481

Pacherie, E. (2007). The Sense of Control and the Sense of Agency. Psyche, 13(1), 1.

Pelli, D. G. (1997). The VideoToolbox software for visual psychophysics: Transforming numbers into movies. Spatial Vision, 10(4), 437–442. https://doi.org/10.1163/156856897X00366

Pereira, M., Skiba, R., Cojan, Y., Vuilleumier, P., & Bègue, I. (2021). Optimal confidence for unaware visuomotor deviations bioRxiv 2021.10.22.465492; doi: https://doi.org/10.1101/2021.10.22.465492

R Core Team (2020). R: A language and environment for statistical computing. R Foundation for Statistical Computing, Vienna, Austria. URL https://www.R-project.org/.

Seow, T., & Fleming, S. M. (2019). Perceptual sensitivity is modulated by what others can see. Attention, Perception, & Psychophysics, 81(6), 1979–1990. https://doi.org/10.3758/s13414-019-01724-5

Sherman, M. T., Seth, A. K., Barrett, A. B., & Kanai, R. (2015). Prior expectations facilitate metacognition for perceptual decision. Consciousness and Cognition, 35, 53–65. https://doi.org/10.1016/j.concog.2015.04.015

Sinanaj, I., Cojan, Y., & Vuilleumier, P. (2015). Inter-individual variability in metacognitive ability for visuomotor performance and underlying brain structures. Consciousness and Cognition, 36, 327–337. https://doi.org/10.1016/j.concog.2015.07.012

Singmann, H., Bolker, B., Westfall, J., Aust, F., & Ben-Shachar, M. S. (2021). afex: Analysis of Factorial Experiments. R package version 1.0-1. https://CRAN.R-project.org/package=afex

Slachevsky, A., Pillon, B., Fourneret, P., Pradat-Diehl, P., Jeannerod, M., & Dubois, B. (2001). Preserved Adjustment but Impaired Awareness in a Sensory-Motor Conflict following Prefrontal Lesions. Journal of Cognitive Neuroscience, 13(3), 332–340. https://doi.org/10.1162/08989290151137386

Sternad, D., Abe, M. O., Hu, X., & Müller, H. (2011). Neuromotor Noise, Error Tolerance and Velocity-Dependent Costs in Skilled Performance. PLOS Computational Biology, 7(9), e1002159. https://doi.org/10.1371/journal.pcbi.1002159

Synofzik, M., Vosgerau, G., & Newen, A. (2008). I move, therefore I am: A new theoretical framework to investigate agency and ownership. Consciousness and Cognition, 17(2), 411–424. https://doi.org/10.1016/j.concog.2008.03.008

Villa, R., Tidoni, E., Porciello, G., & Aglioti, S. M. (2018). Violation of expectations about movement and goal achievement leads to Sense of Agency reduction. Experimental Brain Research, 236(7), 2123–2135. https://doi.org/10.1007/s00221-018-5286-3

Wegner, D. M., & Wheatley, T. (1999). Apparent mental causation: Sources of the experience of will. American Psychologist, 54(7), 480–492. https://doi.org/10.1037/0003-066X.54.7.480

Wolpe, N., Wolpert, D. M., & Rowe, J. B. (2014). Seeing what you want to see: Priors for one’s own actions represent exaggerated expectations of success. Frontiers in Behavioral Neuroscience, 8, 232. https://doi.org/10.3389/fnbeh.2014.00232

Wolpert, D., Ghahramani, Z., & Jordan, M. (1995). An internal model for sensorimotor integration. Science, 269(5232), 1880–1882. https://doi.org/10.1126/science.7569931

Wolpert, D. M., & Flanagan, J. R. (2001). Motor prediction. Current Biology, 11(18), R729–R732. https://doi.org/10.1016/S0960-9822(01)00432-8

Yon, D. (2020). Enhanced metacognition for unexpected action outcomes. PsyArXiv. https://doi.org/10.31234/osf.io/3vn96

